# Unsupervised, frequent and remote: a novel platform for personalised digital phenotyping of spatial working memory and image recognition in humans

**DOI:** 10.1101/2022.08.24.505107

**Authors:** Marius Bauza, Marino Krstulovic, Julija Krupic

## Abstract

Spatial working memory and image recognition tests are commonly used to facilitate the diagnosis of hippocampal-related neurological disorders such as Alzheimer’s disease due to their relatively high specificity and sensitivity to damage to the medial temporal lobes compared to standard commonly used clinical tests. Pathological changes in Alzheimer’s disease start years before the formal diagnosis is made, partially due to testing too late. To address this challenge, we developed a novel digital platform, hAge (‘healthy Age’), which integrates double spatial alternation, image recognition and visuospatial tasks for frequent remote unsupervised assessment of spatial and non-spatial working memory. 191 healthy adults (67% females, 18-81 years old) participated in the study. In line with findings using standard laboratory tests, we showed that performance on the spatial alternation task negatively correlated with inter-trial periods and performance levels on image recognition and visuospatial tasks can be controlled by varying image similarity. Importantly, we demonstrated that frequent engagement with the double spatial alternation task leads to a strong practice effect, previously identified as a potential measure of cognitive decline in MCI patients. Finally, we discuss how lifestyle and motivation confounds may present a serious challenge for cognitive assessment in real-world uncontrolled environments.

## Introduction

Neuropsychological tests provide a powerful tool for facilitating the diagnosis of Alzheimer’s disease (AD) and other forms of dementia[1–3]. At the early stage of AD, the pathology is almost exclusively confined to the medial temporal lobe regions (MTL)[4], which is crucial for episodic[5,6] and spatial working memory[6,7], image recognition[6,8] and navigation[9,10]. Hence, neuropsychological tests targeting these functions were shown to be especially suitable for early AD detection and its differentiation from other neurological conditions such as depression[1,2]. Such tests are normally applied every 8-24 months in research laboratories and clinics by a trained specialist to monitor any changes in the condition[1,3]. As a result, the implementation of such tests is limited in scale, frequency, and duration, which was especially exacerbated during the COVID-19 pandemic[11]. This creates an urgent need to introduce new approaches, implementable at scale and frequency. Here, we present a novel digital platform called hAge (‘healthy Age’) for a fully unsupervised remote and frequent long-term assessment of spatial working memory and image recognition in real-world ethologically-relevant environments (i.e. Participants’ homes, workplaces, etc.). hAge platform is based on standard MTL-specific tests executed in a game-like form, which involves frequent few-seconds-long interactions throughout the day. hAge includes image recognition, visuospatial and spatial alternation tasks known to be highly sensitive to damage in the hippocampal[6,7] and prefrontal areas[7,12], both compromised in the early and middle stages of AD[4]. It has been suggested that image recognition may be sensitive to either hippocampal formation or extra-hippocampal MTL regions (perirhinal, lateral entorhinal and parahippocampal cortices), depending on the strategy an individual uses to solve the task. The use of a cognitively less demanding familiarity strategy has been proposed to rely on extra-hippocampal MTL, while the use of the cognitively more demanding recall strategy is thought to depend on the hippocampus[8,13], but see[14]. Visuospatial working memory task has been shown to depend on both the hippocampus and the prefrontal areas[7].

To demonstrate the feasibility of our approach, we tested whether we could achieve sufficient levels of adherence and whether the performance on hAge tasks is comparable to the performance observed in the analogous standard tests measured in the controlled laboratory environments. Based on previous studies in humans and rodents[7,15], we expected performance on the spatial alternation task to decrease with longer time intervals between trials. Furthermore, we predicted that performance could be modulated by adjusting the difficulty of the tasks, e.g., selecting more similar images or introducing more complex alternation rules. We also expected to see improvements in performance with repeated exposures to the test known as practice effects[16–18]. Finally, we expected to find that performance negatively correlates with age[19,20]. All of these measures were previously used to assess cognitive decline and the risk of developing AD in MCI patients.

## Methods

### Participants

This study has been approved by the University of Cambridge Psychology Research Ethics Committee (PRE.2020.053). Written informed consent was obtained from all study Participants, and the data were analysed anonymously. Participants were recruited via the Cambridge Psychology Research sign-up system and the Join Dementia Research database. Volunteers were included in the study if they declared that they met the following criteria: (1) above 18 years old; (2) had no visual or hearing impairments; (3) had no learning disabilities or dementia; (4) had no history of neurological psychiatric disorders; (5) had access to a Windows PC. Engagement rules were provided in the Participant Information Sheet and the Participant Consent Form and clarified via email, phone call or Zoom session upon request. Participants were explicitly asked to participate in the study for eight weeks and were offered a small £10/week reward if they engaged with the program on average >15 times/day, paid at the end of each week via bank transfer or cheque.

The program was run in two separate 8-week phases (Phase 1 and Phase 2). The participation data were stored on local user computers. The data was encrypted and uploaded to a secure University of Cambridge website several times a day. After the study was concluded, data was anonymised by deleting the table linking the Participants’ identities and aliases.

### hAge tasks and design

The hAge program was designed to simultaneously test three different types of medial temporal lobe dependent tasks over a long period of time in a completely unsupervised way. The first set of tasks was variants of a spatial alternation task (Fig 1a, a yellow inset). The second set involved either image recognition or image-in-place (also known as visuospatial) recognition tasks (Fig 1a, a red inset). Both sets were run simultaneously on Participants’ computers. The hAge program (research.pdn.cam.ac.uk/hkage) was custom-written in Python for Windows OS.

**Fig 1.**
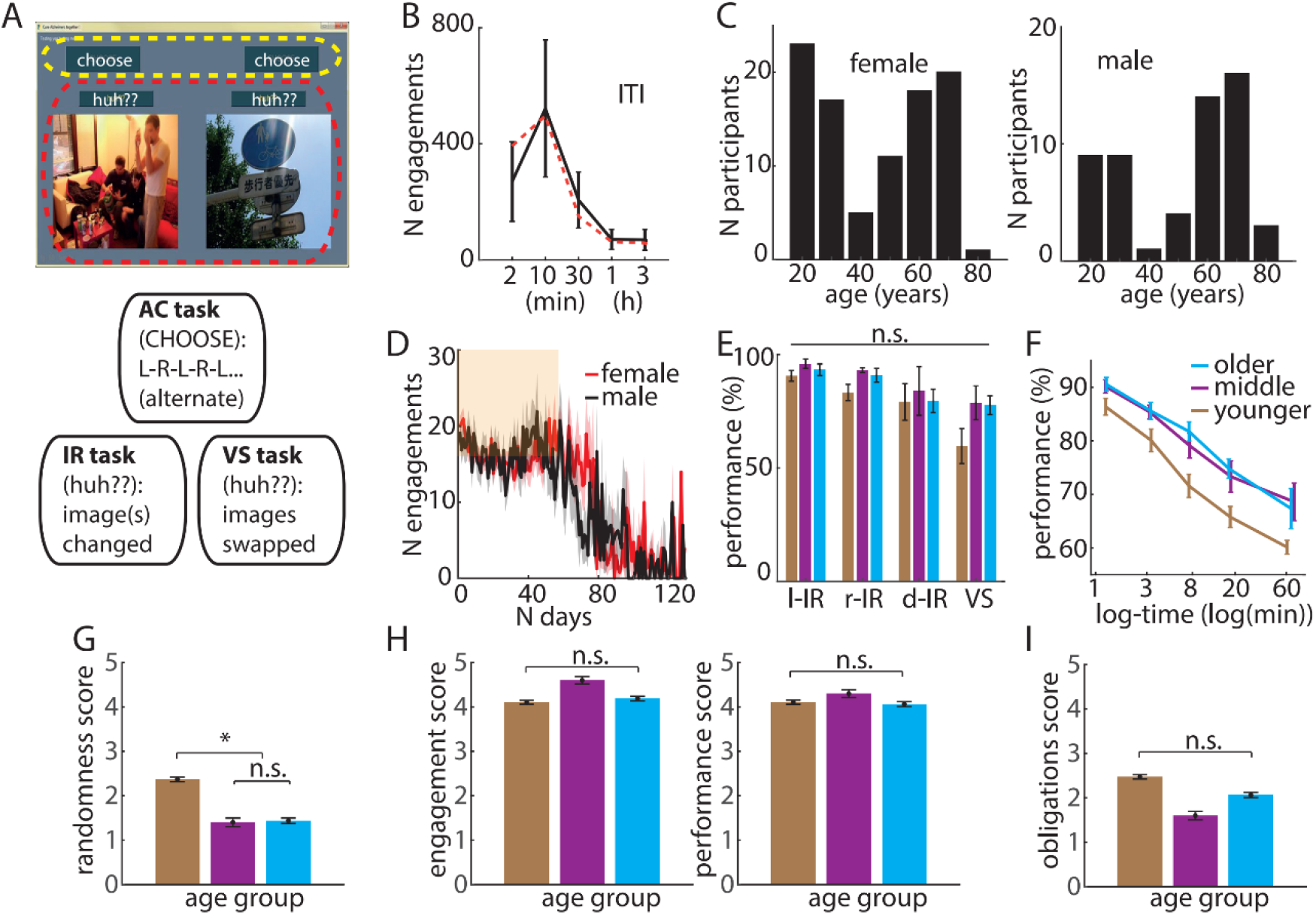
Phase 1 testing of hAge feasibility. (A) Phase 1 hAge task design. The program consists of three types of tasks: 1) spatial alternation (SA), during which Participants clicked one of the **[choose]** buttons (yellow inset), alternating between left (L) and right (R) on every engagement; 2) image recognition (IR) and 3) visuospatial (VS) task (red inset): Participants clicked **[huh??]** button above the image that was perceived to have changed (left or right IR, l-IR and r-IR, respectively). Both left and right **[huh??]** buttons were clicked when both images changed (d-IR) or swapped sides (VS). (B) Most of the time the Participants engaged with the program shortly after it became active. Black solid line indicates the actual distribution of inter-trial intervals (ITIs±s.d.). Red dashed line indicates the hard-coded ITI distribution at which the hAge window became active. (C) Female (left) and male (right) distribution by age. (D) Daily engagement levels throughout the study. Zone of rewarded participation (8 weeks) is shown in orange. Red and black mark female and male engagement levels respectively. (E) All age groups showed high performance on IR and VS tasks. (F) Performance on the SA task was negatively correlated with the inter-trial period in all age groups. (G) The average randomness score showing how often Participants clicked a random button to complete their engagement with the program. 1 to 5 =‘never’ to ‘very often’. (H) The average engagement score (left) showing how motivated the Participants were to achieve the daily required engagement quota; and the average performance score showing how motivated the Participants were to do well when providing their answers. 1 to 5 = ‘not motivated’ to ‘extremely motivated’. (I) The average obligations’ score showing how overwhelmed the Participants were with other professional and/or social/personal obligations. 1 to 5 = ‘not at all’ to ‘extremely overwhelming’. Different colours represent different age groups and are maintained throughout the figure.

The hAge program was tested in two phases (Phase 1 and Phase 2). We used Phase 1 as a proof-of-principle pilot experiment to test the feasibility of our approach. To this end, we implemented the simplest versions of the tasks: images in the image recognition and visuospatial tasks were drawn from completely different categories (e.g. people vs highways, etc.), and the rule of the special alternation task was to alternate between consecutive choices in order to make a correct choice.

In Phase 2, we made some alterations to the program code to fix technical glitches reported by the Participants. Based on their feedback, we also altered the program’s activation sound (see below). Finally, all tasks were made substantially harder to provide a greater spread of the performance scores. Namely, simultaneously displayed images were drawn from the same category (planes, dogs, highways or buildings), whereas a single alternation task was changed to a double spatial alternation task, which required the Participant to alternate sides on every second choice. In both Phases, Participant was immediately provided with feedback on whether their choice was correct on alternation tasks but not on image recognition or visuospatial tasks.

The hAge interface comprised two images and two pairs of buttons, labelled **[choose]** and **[huh??]**, displayed in the same grey window presented at random time intervals (Fig 1A). The time interval between the window appearances (inter-trial interval) varied from ∼25 seconds to ∼3 hours (Fig 1B). The program window was invisible at all other times to prevent any type of ‘rehearsal’. A gentle repetitive sound was played to signal that the program was activated. The program window remained active until the engagement was completed (i.e. Participants indicated their choices). The Participant could also ignore the prompting and re-engage at their own time. During the engagement period, the Participants were presented with both the spatial alternation and the image recognition tasks within the same window (Fig 1A, yellow and red insets, respectively). In Phase 1, to make a correct choice on the spatial alternation, Participants had to alternate their responses between the trials by clicking the left or right **[choose]** buttons. In Phase 2, the Participant had to alternate on every second choice. The concurrently presented image recognition task required Participants to click one or both of the **[huh??]** buttons if they thought that the image beneath the button had changed since the last engagement. If both images changed or swapped places, the Participants were required to click both **[huh??]** buttons. Participants were provided with instructions on how to conduct the tasks prior to commencing their engagements.

### Inclusion criteria

To provide a reliable longitudinal assessment of spatial working memory, we only included data from Participants who had engaged with the spatial alternation task for at least 6 out of 8 weeks with a daily engagement of ≥10/day. The duration and daily engagement were decided prior to the data collection. The image recognition or visuospatial tasks were included only when the former criterion was met and when the total number of engagements in the image recognition or visuospatial tasks was >10 times in total (for each of these tasks). To assess learning (practice) effects on the image recognition tasks, the performances were split into two equal blocks (early vs late) and were included in the final analysis if there were at least 10 measurements on each task per block. During Phase 2 frequency of image changes was increased from ∼1 change/day (Phase 1) to ∼5 changes/day (Phase 2) to assess learning effects.

We also excluded a small number of Participants (n=17, S1 Table) who showed abnormally high (average performance >90% across all ITIs) or unusually low performance levels (<60%) under the assumption that they likely misunderstood the task by using external reminders (high performers, n=7) or choosing mostly randomly (low performers, n=10). It must be noted that the main results do not change if we do not exclude these Outliers.

### Spatial Alternation and double Spatial Alternation Tasks

The performance on the alternation tasks was evaluated at five different inter-trial intervals (ITI): (1) <2min; (2) 2-5 min; (3) 5-10 min; (4) 10-30 min, and (5) >30min to ensure adequate sampling for each interval (>100 samples per interval). The median of all values within a respective bin is shown as the position of that bin in plots. The ITIs are displayed as log(min) to facilitate data visualisation.

The performance on the double alternation task was divided into win-shift (Participants had to alternate a previously chosen side to make a correct choice) and win-stay task (Participants had to hold on to a previously chosen side to make a correct choice).

### Statistical Analysis

The data was analysed using MATLAB (The MathWorks Inc.). All statistical tests are stated in the main text and Supplementary Methods. Paired-sample t-test was used to estimate if there was a significant difference in performance on the single and double alternation tasks and between the win-stay and win-shift components of the double alternation task. All ITIs were included separately. One sample t-test was used to estimate if there was a significant difference between IR and VS tasks in different Phases. One-way ANOVA with Bonferroni correction was used to estimate the effect significance of different age groups. Single and double image recognition tasks were combined for statistical analysis. Kruskal-Wallis H test was used to compare the questionnaire results between different age groups.

The chance level for image recognition and visuospatial tasks was calculated by assuming that upon reactivation of the program window, a Participant would randomly select that one or both images changed, with an equal probability between all four visual tasks.

### Correlation Analysis

Spearman’s rank correlation coefficient was used to estimate the change in performance on double spatial alternation task over days.

### Feedback Collection

At the end of the eight weeks, Participants were informed that their participation was completed and they would no longer receive any payment, although they could continue using hAge if they wished to, and the collected data would be used in the study. We also asked Participants to complete a short feedback form and a demographics questionnaire (S1 Fig). Phase 2 Participants, who passed the inclusion criteria, were additionally asked to complete a short follow-up questionnaire regarding their lifestyle and motivation while they used the hAge program (S1 File) and were invited to participate in the Focus Group discussion (S2 File) to help better understand their experience of performing the tasks and the strategies they used to solve them. The questionnaire consisted of 5 questions which were on a 5-point Likert scale. Out of the 72 people that were emailed, 45 (62.5%) responded to the questionnaire. Of the 45 respondents, 16 were from the older, 10 from the middle and 19 from the younger age groups. 10 Participants were selected for the follow-up Focus groups. Each Focus Group comprised 3-4 Participants, and two researchers involved in the study, and the meeting lasted ∼1 hour. Altogether, we conducted 3 Focus group meetings.

## Results

### Phase 1: demonstrating the feasibility of hAge for longitudinal, remote and unsupervised assessment of spatial working memory and image recognition

In total, 151 healthy adults (18-81 years old; mean±s.d.: 48±20 years old; 63% females, Fig 1C, S1 Fig, S2 Table) took part, of whom 22% (67% females) passed the inclusion criteria (S2 Fig; engaged with the App for at least 6 weeks, see Methods) and were used for subsequent analysis. It must be noted that the majority of Participants were excluded due to ‘insufficient’ number of days they engaged with the program (S2 Fig; less than 42 out of 56 days; with the majority of Participants engaging for at least ∼30 days) and not due to the low number of daily engagements. The age distribution was comparable between these two populations. On the other hand, we also found that many Participants continued to play beyond the requested 8-week period (Fig 1D, S2 Fig) at high daily engagement levels, with many reporting that they liked using the program and wished to continue playing (S3 Table), supporting the feasibility of such active frequent unsupervised testing.

All age groups showed high performance on image recognition and visuospatial tasks with no significant difference between the groups (Fig 1E; combined IR task: F(2,72)=1.01, P=0.37; and VS task: F(2,26)=2.33, P=0.12, one-way ANOVA test). The majority of errors on the visuospatial task resulted from missing the change.

In line with our initial hypothesis, we also found that performance on the spatial alternation task rapidly decreased with longer ITIs (Fig 1F). However, unexpectedly, the average combined performance was lowest in the younger group, with no differences between the middle and older groups (Fig 1F, mean±s.m.e.: 76.0±1.4%, 82.0±1.6%; 83.2±1.2% in younger, middle and older age groups respectively; P=0.0005, F(2,129)=8.12, one-way ANOVA) even though the performance on analogous clinic-based working memory tests typically negatively correlates with age[21]. Based on previous studies[22,23] and the results from the follow-up Questionnaire (S1 File), higher absolute performance of the middle and older groups may be attributed to differences in commitment levels, with older and middle groups likely dedicating more cognitive effort in an attempt to correctly complete the task. Namely, when the Participants were asked if they often made a random choice simply to end the engagement and return to their previous activities, the younger group did this significantly more often compared to older and middle age groups (Fig 1G, randomness score mean±s.m.e.: 2.4±0.3, 1.4±0.2; 1.4±0.2 in younger, middle and older age groups respectively; P = 0.01, X^2^ (2, 42) = 9.2, Kruskal-Wallis H test). Interestingly, there was no significant difference between the groups in the subjective perception of their commitment and motivation (Fig 1H). In addition, the younger group tended to have higher levels of perceived distraction, possibly due to more active lifestyles, although the difference was not significant (Fig 1I, obligations’ score mean±s.m.e.: 2.5±0.2, 1.6±0.3; 2.1±0.2 in younger, middle and older age groups respectively; P=0.099, X^2^(2, 42)=4.63, Kruskal-Wallis H test). The assumption of lower commitment levels in a younger group was further corroborated by a precipitous decline in average performance of a younger group on the spatial alternation task, which happened exactly after the required 8 weeks and which was not observed in the older and middle groups (S3 Fig).

### Phase 2: evaluating performance on more challenging tasks

The second, more challenging version (Phase 2) of the hAge program with modified SA, VS and IR tasks increased the cognitive demand required to solve them. Based on findings from analogous standard tests, we hypothesised that the Participants would show reduced performance on all tasks. This is important because the high average performance observed during Phase 1 may lead to a ceiling effect and limit the usability of our approach. Instead of a simple SA task, we implemented a double spatial alternation (dSA) rule (see Methods). Furthermore, co-displayed images in IR and VS tasks were drawn from the same image category, making them significantly more similar and hence more challenging to spot the change (Fig 2A). In total 72 adults (19-73 years-old; mean±s.d.: 53±15 years-old; 67% females) participated in the Phase 2 release (Fig 2B), of whom 54% passed the inclusion criteria (22-73 years old, 74% females; Fig 2C-D). 44% (32/72) also took part in Phase 1 to allow direct comparison. Importantly, of the 40 new Participants taking part in Phase 2, 65% (26/40) passed the inclusion criteria suggesting that modifications introduced in Phase 2 (fixing technical glitches, changing the activation sounds and modifying the cognitive tasks) were able to significantly improve the adherence levels compared to Phase 1.

**Fig 2.**
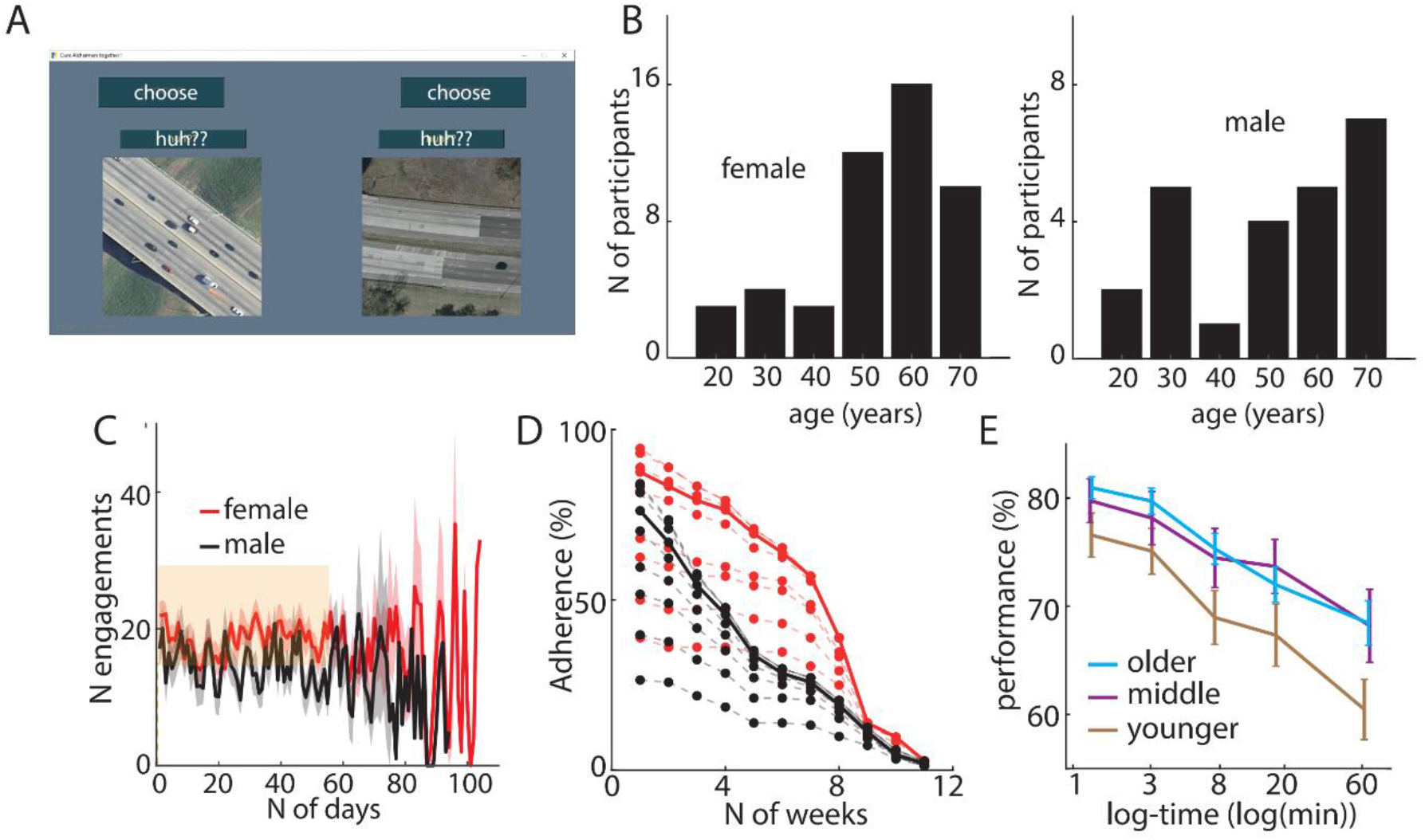
Performance on Phase 2 hAge program. (A) Phase 2 hAge was significantly more challenging compared to Phase 1. The SA task was changed to a double spatial alternation (dSA) task where Participants had to alternate on every second choice to pick the correct side (LLRRLLRR). Simultaneously presented images on IR and VS tasks were drawn from the same categories. (B) Female (left) and male (right) distribution by age. (C) Daily engagement levels throughout the study. Zone of rewarded participation is shown in orange. (D) The adherence levels at different durations and daily participation in Phase 1 (black) and Phase 2 (red) programs. The x-axis corresponds to the number of weeks of engagement. Solid lines show the minimum daily engagement level equal to 10 used in the analysis. Dashed lines correspond to minimum daily engagement levels set to 4, 6, 8 (above solid lines) and 12, 14, 16, 18, 20 (below solid lines). As expected, the adherence levels fall with the increased daily engagement and the total required duration of the engagement. (E) As in the Phase 1 SA task, the performance on the dSA task was negatively correlated with the inter-trial period in all age groups.

As expected, the Participants performed significantly worse on dSA compared to the SA task (mean difference ± s.m.e.: 8.4±0.5%; P = 4.3 × 10^−33^, t=-15.4, df=155 paired-sample t-test), likely due to increased memory load requiring to remember the last choice and the order within the choice sequence. Similar to Phase 1, the performance on the dSA task declined with increasing ITIs, and the younger group still showed the lowest average performance with no significant difference between the middle and the older groups (Fig 2E; P=0.0033, F(2, 153)=5.93, one-way ANOVA). Interestingly, performance on the dSA task varied depending on the type of choice Participants were making. Namely, in one case, the correct choice was the opposite from the previously selected one, requiring to alternate (win-shift), whereas in another case, the Participant had to repeat the previous choice (win-stay). The average performance on the win-shift task was significantly lower compared to the win-stay task (Fig 3A, mean±s.m.e.: 72.9±0.7% vs 78.3±0.9%, P=1.1 × 10^−6^, t = -4.97, df = 310, two-sample t-test), suggesting that Participants found it more difficult to remember to alternate compared to holding on to the same side. Surprisingly differences in performance on win-shift and win-stay tasks (the relative performance) in the older group were significantly larger compared to younger and middle age groups, with no difference between the latter two (Fig 3B, mean±s.m.e.: -9.0±1.3%, -3.8±1.4%; -3.3±1.4% in older, middle and younger age groups respectively; P=0.0055, F(2,153)=5.38, one-way ANOVA).

**Fig 3.**
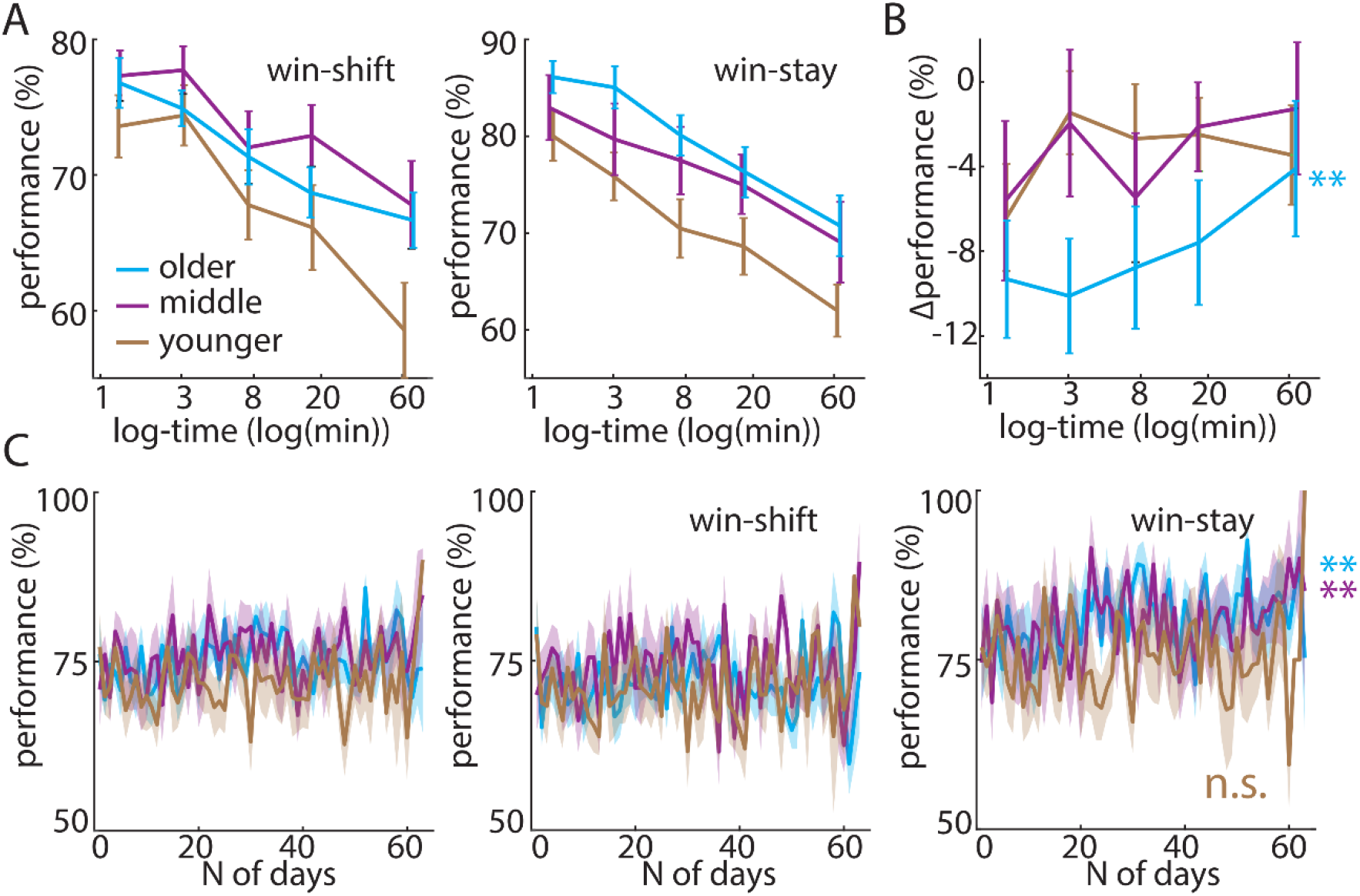
Different performance on win-shift and win-stay dSA task. (A) All age groups showed different performance on win-shift (alternate the choice) vs win-stay (keep the same choice) dSA tasks. (B) The relative performance (difference between performance on win-shift and win-stay tasks) was significantly larger in the older group compared to middle and younger groups with no difference between the latter ones. (C) Overall performance (left) did not change with experience in any of the age groups. However, while the performance on the win-shift task requiring remembering alternate choices did not improve with experience, the performance on the win-stay task (i.e. remembering to hold on to the same choice) kept on significantly improving in the older and middle groups but not in the younger group.

We next investigated whether our longitudinal assessment allowed us to detect any practice effects previously used to evaluate cognitive decline in MCI patients[16–18]. To address this question, we looked whether there were any changes in win-shift and win-stay task performance over time. Indeed, the performance of older and the middle groups on the win-stay task improved significantly with experience (Fig 3C, ρ=0.40, P=0.0013; and ρ=0.36, P=0.004, respectively; Spearman’s Rank-Order Correlation) while the performance on the win-shift task on average remained similar over time (ρ =-0.03, P=0.80 and ρ=0.002, P=0.99, respectively; Spearman’s Rank-Order Correlation). Notably, the performance did not improve with experience in the younger group on either win-stay or win-shift tasks (ρ=0.089, P=0.49 and ρ =0.14, P=0.27, respectively; Spearman’s Rank-Order Correlation).

As expected, the overall performance on image recognition and visuospatial tasks (Fig 4A) was significantly lower in Phase 2 compared to Phase 1 (Fig 4B; mean difference ± s.m.e.: 34.9±2.8%, P=1.6 × 10^−21^, t=-12.6, df=90 one-sample t-test). The new task provided a wide spread of scores, avoiding both ceiling (100% performance) and floor (chance level ∼5%, see Methods) effects. Unlike findings in Phase 1, there was a significant age effect with the older group again outperforming other groups (combined IR mean±s.m.e.: 71.4±3.9%, 50.9±4.2% and 53.6±4.2% in the older, middle and younger groups, respectively P=0.0007, F(2, 111)=7.71; and VS mean±s.m.e.: 70.8±7.3%, 52.9±7.9% and 51.8±7.9% in the older, middle and younger groups, respectively P=0.15, F(2, 35)=2.0 one-way ANOVA; N.B. IR and VS tasks showed the same trends but the latter did not reach the significance possibly due to a smaller sample size). In general, given relatively high image similarity, the majority of errors for all age groups occurred when the Participants failed to notice any change, which was more pronounced in the middle and younger groups, while the other types of errors remained similar between the groups (Fig 4C). The majority of such false negatives were made during VS task, while the least of them occurred on the d-IR task, reflecting different difficulty levels of these tasks (Fig 4D). There was no overall noticeable improvement in VS or IR tasks in any age group over time (Fig 4E).

**Fig 4.**
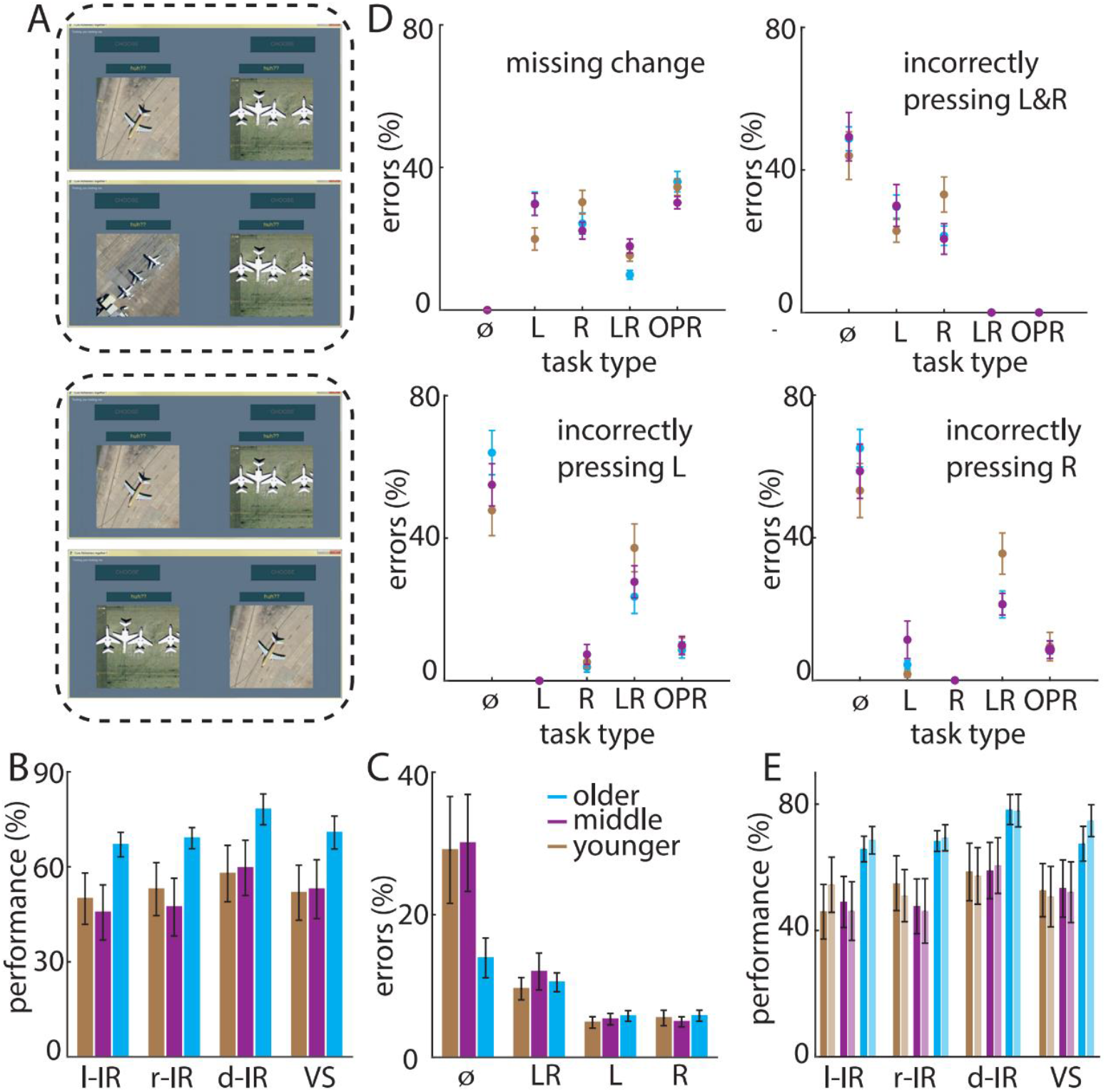
Performance on Phase 2 IR and VS tasks. (A) An example of the left l-IR (top inset) and VS (bottom inset) tasks. All of the images were drawn from the same ‘planes’ category. (B) Performance on Phase 2 IR and VS tasks was lower compared to Phase 1. (C) The type of errors on IR and VS tasks for each age group. (D) Breakdown of choices associated with different types of errors (specified in legend). Task types are shown on the x-axis. ⌀ denotes no image change taking place. All age groups mostly missed the change in VS task (top left) while they tended to overreact to no change (⌀) by incorrectly pressing left (L), right (R) or both (LR) buttons. (E) The performance on different IR and VS tasks did not noticeably improve with experience: the first half of total engagements are shown in dark and the latter half in light colours for each age group.

## Discussion

Here we described a novel digital game-like platform for remote longitudinal unsupervised and frequent assessment of spatial working memory and image recognition known to rely on the hippocampal-transentorhinal circuitry in humans and other mammals[6–8,24–27]. We report a 65% (26/40 of new Participants in Phase 2) Participant adherence level, comparable to other similar state-of-the-art unsupervised platforms[28,29]. The adherence level during Phase 2 was markedly improved from 22% achieved during the first roll-out of the program (Phase 1). We anticipate that the adherence levels will be further improved by implementing our approach on mobile devices.

Our results show that in line with previous findings on analogous spatial memory tests conducted in clinics and research laboratories, the performance on spatial alternation and double spatial alternation tasks negatively correlates with inter-trial intervals. Importantly, unlike in other tests, we could simultaneously sample the whole range (20 seconds – 3 hours) of inter-trial intervals. Furthermore, the task difficulty levels could be dynamically adjusted to optimise the spread of scores and avoid both ceiling and floor effects. Namely, the level of difficulty could be controlled by introducing more complex rules of spatial alternation or by making the images more similar in image recognition and visuospatial task. Notably, we found strong practice effects on double spatial alternation task in older and middle age groups (>50 years old). Previously, improvements on similar spatial memory tasks were identified as potential measures for assessing cognitive decline in MCI patients[16–18].

Unexpectedly, we found that the older and middle age Participants demonstrated higher absolute performance on all hAge tasks compared to younger Participants (<50 years old). This suggests that even though remote testing may solve frequency and scalability limitations associated with standard neurophysiological assessments, it brings an entirely different set of challenges related to testing in uncontrolled environments. We hypothesise that in real-world environments, Participants’ ability to solve a given task is often significantly influenced by other factors not directly related to spatial working memory and image recognition, such as cognitive load, external distractions, motivation and compliance. The analysis of questionnaire data further supported this hypothesis. Given such confounds, we anticipate that such unsupervised active testing may be more suitable for older age groups (>50 years old), who may be motivated by their wish to receive an unbiased assessment of their spatial cognitive abilities, which may have clinical relevance. The interest in clinical relevance in older groups was confirmed during the Focus Group discussions. We speculate that using relative rather than absolute performance measures may be more appropriate in uncontrolled environments. This can be achieved by creating personalised double spatial alternation, image recognition and visuospatial tasks with different degrees of difficulties by adjusting the time intervals between the choices (double spatial alternation task), applying different choice rules or by tuning similarity between the presented images as was done in Phase 1 and Phase 2 image recognition and visuospatial tasks. Such personalised readjustment will impose different cognitive demand required to solve the tasks while keeping other external factors similar.

In summary, our results show that hAge presents a promising new approach for mass testing MTL-related spatial working memory and image recognition under ethological conditions in a remote and unsupervised way. To validate its utility for clinical diagnostics on a large scale, it will have to be tested in clinical populations and implemented on mobile devices instead of desktop PCs. This should dramatically increase its flexibility and allow to link the performance outcomes to external factors routinely measured by mobile devices (e.g. sleep, exercise, location etc.). Moreover, the platform will have to be validated against well-established AD biomarkers and correlated with imaging measures of MTL and prefrontal areas.

## Acknowledgements

We thank all of the Participants for their participation in this study. We also thank Profs. Rik Henson, Dennis Chan and John O’Keefe for their comments on the early version of the manuscript.

## Author Contributions

M.B. and J.K. conceived the study. M.B. developed the hAge program. M.K. recruited the Participants and acted as the Participant liaison over the course of the study. All authors managed the weekly payments. M.B and J.K. analysed the data and wrote the manuscript with contributions from M.K.

## Supplementary Information

is available for this paper.

## Materials & Correspondence

Correspondence and requests for materials should be addressed to J.K. and M.B.

## Code availability

All custom code written for reported analyses will be available via request to the corresponding authors.

## Data availability

The anonymised dataset for all experiments will be publicly available via request to the corresponding authors.

## Supporting information

**Figs. S1-S3**

**Tables S1-S3**

**Files S1-S2**

