## Supplementary Information for "Unsupervised, frequent and remote: a novel platform for personalised digital phenotyping of spatial working memory and image recognition in humans"

**S1 Fig. The highest degree of schooling achieved across all age groups.**

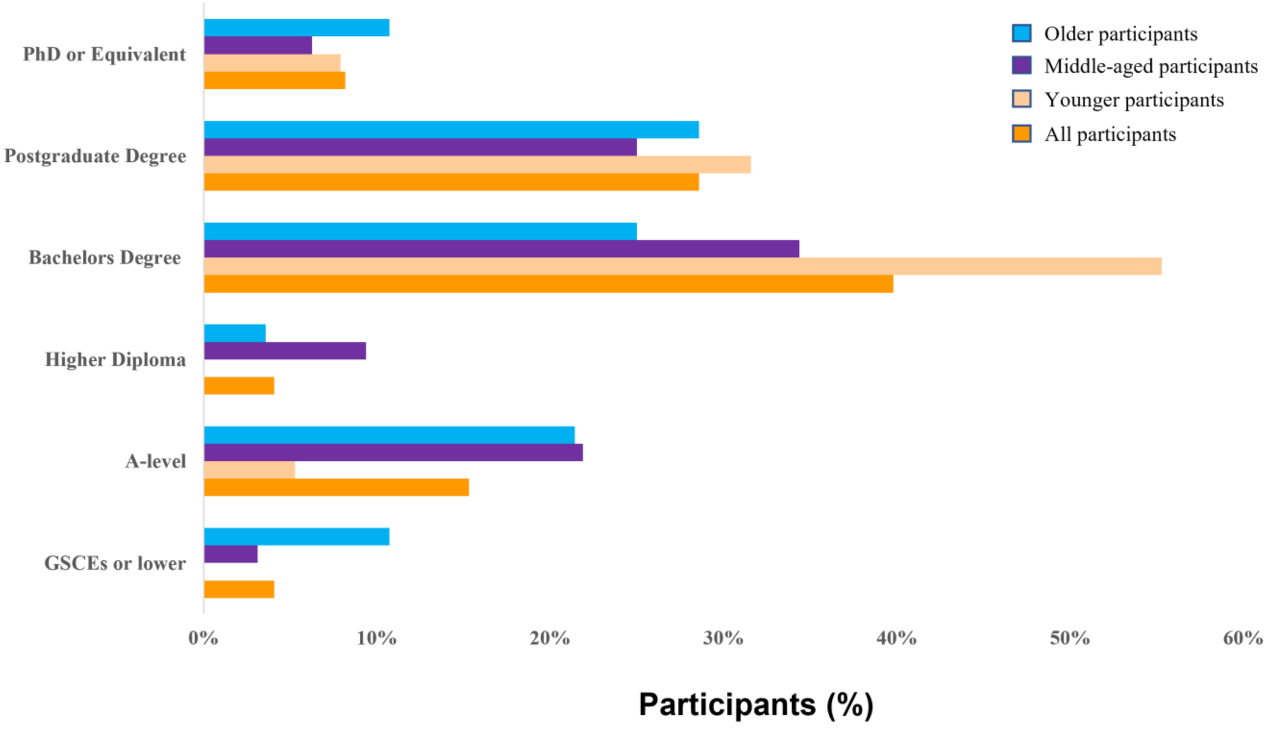

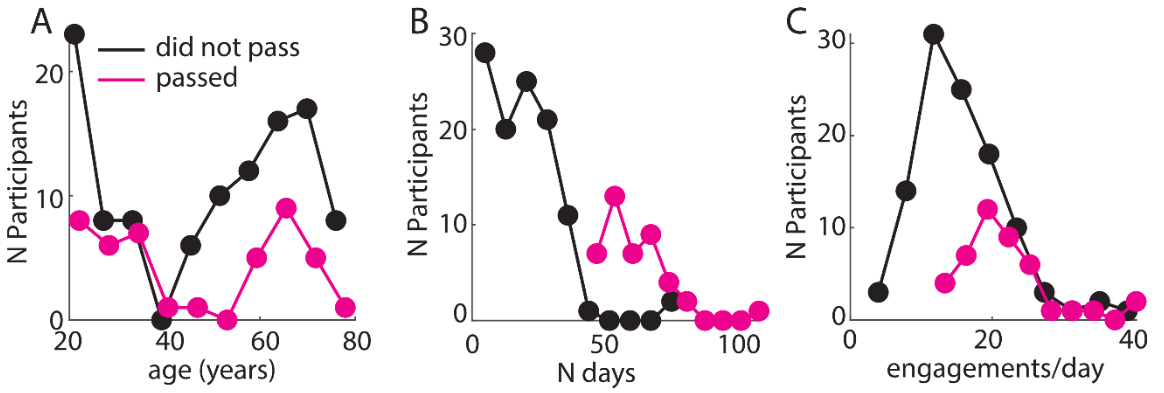


**S2 Fig. Participants' adherence levels during Phase 1.** (A) Age distribution was comparable in Participants who passed inclusion criteria (magenta) and those who did not (black). (B) The number of days of engagement was the major factor determining factor contributing to the exclusion of the Participants with the majority of Participants engaging with the program for ~30 days. (C) The average daily participation levels were sufficiently high in both groups.

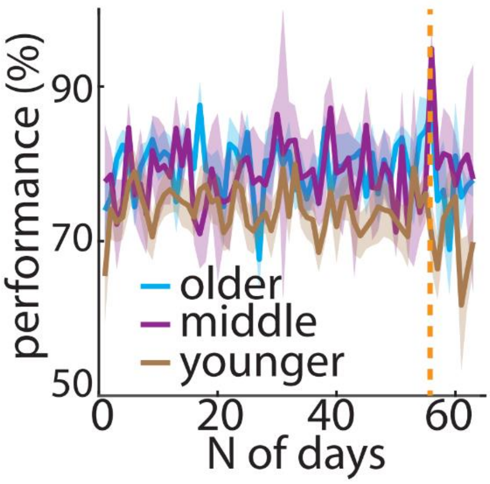


**S3 Fig. Performance on SA task measured over time.** Performance on the SA task did not change with experience in the older and middle age groups while it steeply declined in the younger group right after the end of the study, i.e. requested and paid 8 weeks indicated with the dashed line. Different colours indicate different age groups.

**S1 Table. Distribution of outliers across different Phases and age groups.**

| Phase | Age group | Total number of Outliers | High performers |
|---|---|---|---|
| 1 | younger | 6 | 2 |
| 1 | middle | 1 | 0 |
| 1 | older | 3 | 3 |
| 2 | younger | 2 | 0 |
| 2 | middle | 4 | 1 |
| 2 | older | 1 | 1 |

**S2 Table. Participants' Demographic Information**

| Demographics | All Participants (n = 99) | Older Participants (n = 28) | Middle-aged Participants (n = 32) | Younger Participants (n = 38) |
|---|---|---|---|---|
| Age (years), mean SD | 50.21 (1.37) | 69.25 (3.23) | 58.75 (4.66) | 29 (7.72) |
| Gender female n(%) | 64 (65.31%) | 17 (60.71%) | 21 (65.63%) | 26 (60.71%) |
| Exercise (times per week), mean SD | 3.69 (2.28) | 4.46(2.11) | 4.06(2.07) | 2.81(2.05) |
| Number of hours of sleep per night, mean SD | 6.95 (1) | 6.78(0.91) | 6.62(1.09) | 7.36(0.85) |

**S3 Table. Participants' Feedback**

| | |
|---|---|
| **Participant No 22** | I would like to thank you for the possibility of taking part in this study. It has certainly proved an eye-opener for me when it comes to assessing my memory. |
| **Participant No 25** | I am really enjoying and find that made me less blaze and focus. Before, I had an attitude that, if was of no importance to me, I did not need to remember. All that changed, thanks to you all giving me this opportunity. I hope from now on, I can carry on helping in whatever way that I can. |
| **Participant No 77** | I have downloaded the new app and I am now using it. The revised tasks are an interesting challenge and I can already see traits in my own capability which are quit revealing even if I can't explain them scientifically! I'm happy to continue with the new app for as long as you find it helpful. |
| **Participant No 80** | Thanks for inviting me to participate - I really enjoyed it and like the idea of making a difference.<br><br>I may continue using the app for a little while |
| **Participant No 86** | It has been a pleasure to participate in the study and I am happy to continue to do so if it is of help to you regardless of reward - kind and useful though that has been. |
| **Participant No 90** | Many thanks for your email and for the compensation over the last few months. I hope my stumbling through your app has given you some useful data. I have filled in your questionnaire and attached it to this email. I have also completed the online questionnaire. It's been an interesting personal challenge and an eye opener to say the least. I now know why my 5 year old Granddaughter can beat me at memory card games! |

**S1 File. Motivation follow up questionnaire**

While being a participant in the hAge experiment, did you find the amount of obligations in your life (both professional and/or social) overwhelming?

1 2 3 4 5

Not at all Extremely overwhelming

How motivated were you to achieve the daily required engagement quota with the hAge app?

1 2 3 4 5

Not motivated Extremely Motivated

How motivated were you to do well when providing your answers?

1 2 3 4 5

Not motivated Extremely Motivated

1. When the hAge app would notify you about a trial (would start ringing), how often would you randomly click on an answer to resume your previous activity?

1 2 3 4 5

Rarely Very often

Did the hAge task format (brief tasks throughout the day) influence your decision to participate in our experiment for weeks?

1 2 3 4 5

No influence Strongly Influenced

**S2 File. Focus Group Questions**

1) What motivated you to participate in this study using the hKage App and very importantly how did you stay motivated to continue?

2) And could you tell us a bit more about how well you could integrate App into your daily routines?

2.a) Can you tell us about your experience in using digital technologies such as smart phones, tablets and others?

2.b) How would you integrate our app with these technologies?

3) One of the task ('choose') gave you an instantaneous feedback on your performance whether the other task ('huh??') did not. Does it matter do you think to get a quick feedback on your performance?

4) Did you use any strategies to solve the tasks of the App?

5) Let's talk specifically about what changes you would introduce to the existing App to make it better?

6) How about the rewards? As you recall in the current version that we released the participants received £10 reward for their participation, which some chose to donate to a charity of their choice and others decline to take altogether. So different individuals get the feeling of reward and positive stimulation differently. We would like to hear from you, going forward what would be a good reward system which would keep you engaged, if the monetary reward was not available.

7) We would also like to hear your opinion about the sense of community you got while you used this app. None of you were in any way connected which each other or knew about each other. Do you think, this option would be attractive going forward? Just to tell you an example, during the first stages of the development of our App, I asked my mum and a mum of my partner to play the App; and whenever we met as a family our mums shared their experience which gave them a lot of fun.

8) What is your opinion about self-tracking your performance? Would you trust App-based diagnostics or you would prefer the feedback to be managed by your local GP or other relevant qualified health practitioner?

9) And finally, what kind of problems you would envision with such Apps? E.g. are you concerned with the possible security issues related to your personal data? (and if so what you would like to see implemented to elevate your concerns?)
